# The evolution of constitutively active humoral immune defenses in *Drosophila* populations under high parasite pressure

**DOI:** 10.1101/2023.10.02.560629

**Authors:** Shuyu Olivia Zhou, Ramesh Arunkumar, Shuai Dominique Ding, Alexandre B. Leitão, Francis M. Jiggins

## Abstract

Both constitutive and inducible immune mechanisms are employed by hosts for defense against infection. Constitutive immunity allows for a faster response, but it comes with an associated cost that is always present. This trade-off between speed and fitness costs leads to the theoretical prediction that constitutive immunity will be favored where parasite exposure is frequent. We selected populations of *Drosophila melanogaster* under high parasite pressure from the parasitoid wasp *Leptopilina boulardi.* With RNA sequencing, we found the evolution of resistance in these populations was associated with them developing constitutively active humoral immunity, mediated by the larval fat body. Furthermore, these evolved populations were also able to induce gene expression in response to infection to a greater level, which indicates an overall more activated humoral immune response to parasitization. The anti-parasitoid immune response also relies on the JAK/STAT signaling pathway being activated in muscles following infection, and this induced response was only seen in populations that had evolved under high parasite pressure. We found that the cytokine Upd3, which induces this JAK/STAT response, is being expressed by immature lamellocytes. Furthermore, these immune cells became constitutively present when populations evolved resistance, potentially explaining why they gained the ability to activate JAK/STAT signaling. Thus, under intense parasitism, populations evolved resistance by increasing both constitutive and induced immune defenses, and there is likely an interplay between these two forms of immunity.

**Author Summary:** Immune defenses can be induced after infection or they may be constitutively active, even in uninfected individuals. As constitutive immunity is a more rapid response, theory predicts that it will be favored when animals frequently encounter parasites. When we subjected populations of *Drosophila melanogaster* to high rates of parasitization from its natural parasite, *Leptopilina boulardi* parasitoid wasps, we indeed observed that the immune response became constitutively active. Uninfected insects had an activated humoral immune response and produced cytokine-secreting immune cells that were normally induced after infection. However, we also found that these populations evolved a greater induced response. This included a greatly increased cytokine response after infection, suggesting that the constitutive activation of some aspects of the immune system may allow a greater induced response in other tissues.

## Introduction

The innate immune system utilizes both constitutive and induced mechanisms for defense [1–3]. Constitutive immunity is always active, regardless of the presence of infection, whereas induced immune mechanisms are activated only in response to infection. Constitutive defense includes the production of antimicrobial peptides (AMPs) [4] and the presence of circulating immune cells in the absence of infection. On the other hand, induced immunity has the potential to be amplified many times, such as the massive upregulation of AMPs in response to microbial infections and the proliferation and differentiation of immune cells [4]. Constitutive immunity can provide an immediate response and eliminate pathogens in the early stages of an infection. Nevertheless, immunity is costly to organisms, and constitutive immunity diverts energy from other components of fitness to defense [5–7]. While costs from constitutive immunity are always present, inducible defense mechanisms impose minimal costs when pathogens are absent. However, mounting an inducible response can be time-consuming, leading to a trade-off between the speed of response and the fitness costs [2,8].

Theoretical models predict that a key factor for determining whether constitutive or induced mechanisms are favored is the probability of encountering a parasite, where frequent parasite exposure leads to the favoring of constitutive defenses [1,2,9]. Consistent with these models, when we artificially evolved *Drosophila melanogaster* populations under high rates of parasitism by the parasitoid wasp species *Leptopilina boulardi*, we found that the populations evolved a constitutively active cellular immune defense. Immune cells in *D. melanogaster* are called hemocytes and have similar functions to leukocytes in vertebrates. Using single-cell RNA sequencing, we found that immature lamellocytes, a type of hemocyte that appears through differentiation in response to parasitization, became constitutively present in these evolved populations [8]. The total and circulating hemocyte numbers also increased with frequent parasite exposure, again mimicking the induced immune response [8,10]. Moreover, the transcriptional signature of hemocytes from uninfected larvae of these populations also displays constitutive upregulation of immune-inducible genes [8]. All these constitutive defense mechanisms likely contributed to the increased level of resistance in these populations.

When a parasitoid wasp lays its egg inside *D. melanogaster*, the fly larva launches an anti-parasitoid immune response following infection, which can be divided into cellular and humoral responses [4]. Cellular responses include the proliferation of hemocytes and the differentiation of specialist lamellocytes that function to encapsulate the parasitoid egg in a multilayered cellular capsule [11–13]. The capsule becomes melanized through prophenoloxidase activities mediated by crystal cells and lamellocytes, eventually killing the parasite [14]. Humoral immunity is mediated by the fat body of the *D. melanogaster* larva, which is the metabolic hub of the organism, functionally similar to the mammalian liver. The larval fat body is responsible for the secretion of systemic humoral immune effectors. For example, antimicrobial humoral immunity in *D. melanogaster* is mediated by the two hallmark NF-κB signaling pathways in the fat body – Toll and Imd pathways – which upregulate effectors such as antimicrobial peptides [4]. However, the role of humoral immune mechanisms in anti-parasitoid immunity is not well characterized. A few secreted immune effectors have been shown to be involved in this response, including thioester-containing proteins (TEPs), where their mammalian counterparts are complement factors involved in the complement cascade [15], a C-type lectin called *lectin-24A* shown to be crucial in the encapsulation response [16], and serine proteases which play key roles in the melanization reaction and Toll pathway activation [17,18].

In addition to the involvement of the two immune tissues, fat body and hemocytes, in the anti-parasitoid response, there exists an interplay between hemocytes and somatic muscles in the *D. melanogaster* larvae. Wasp infection induces the expression of the cytokines Upd2 and Upd3 by circulating hemocytes, which induce of JAK/STAT activity in somatic muscles [19]. Furthermore, JAK/STAT activation in muscles is shown to be required for the encapsulation response, including lamellocyte formation [19].

In this study, we investigated how the constitutive and induced humoral immune defenses have evolved with high rates of parasitism. We conducted RNA sequencing on the larval fat bodies of *D. melanogaster* populations evolved under high parasite pressure or no parasite pressure. This allowed us to examine the expression of immunity genes changed as resistance evolved, and whether these changes were constitutively present before infection or were inducible after infection.

## Results

### The evolution of resistance leads to constitutive expression of immune-inducible genes in the larval fat body

Six experimental populations of *D. melanogaster* were established from 377 wild-caught *D. melanogaster* females [8]. Three of these (N1-3) were maintained with infection by the NSRef strain of the parasitoid wasp *Leptopilina boulardi* at every generation, where flies that survived by launching a successful encapsulation and melanization response were used to establish the next generation [8] (Figure 1A). The other three populations (C1-3) were not parasitized but were otherwise maintained under the same conditions. The populations maintained with high parasitism evolved resistance. After 56 generations, about 50% of flies successfully encapsulated and melanized the wasps, while the no parasitism controls were only able to melanize less than 10% (Figure 1B; Welch’s *t*-test, d.f. = 2.09, *p*-value = 0.0064).

**Figure 1.**
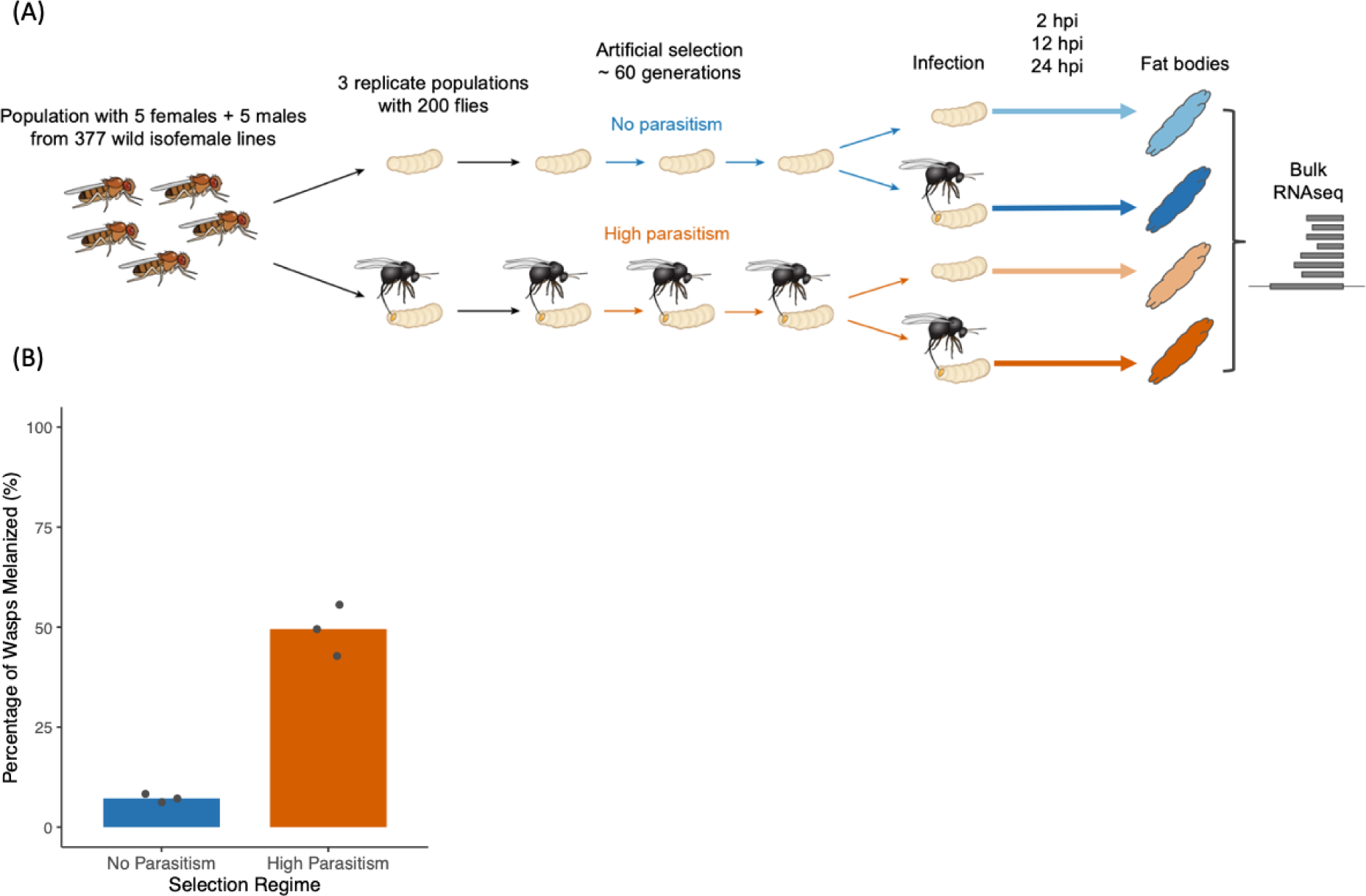
Selection of *D. melanogaster* populations for resistance to parasitoid wasp *Leptopilina boulardi*. (A) Schematic of experimental design. RNA sequencing was conducted at around the 60th generation of selection. (B) The rate of successful melanization of parasitoids in populations maintained with no parasitism and those maintained with high parasitism after 56 generations. Bars represent the mean per selection regime. Points represent replicate populations. Ten replicates of around 50 larvae were assayed per population.

We found previously that high parasitism rates had led to a more active cellular encapsulation response in the populations, including the constitutive presence of immature lamellocytes [8]. A constitutive increase in the number of circulating hemocytes was also observed in these populations [8]. To investigate how humoral immunity has evolved in *D. melanogaster* populations under high pressure of parasitism, we collected the fat bodies of male larvae by dissection from each of the six populations. We analyzed the transcriptional response to infection at 2-, 12-, and 24-hours post-infection (hpi), as well as the transcriptome of fat bodies from age-matched uninfected larvae (Figure 1A).

We investigated whether the humoral immunity of the selected populations would exhibit a similar constitutively active state as the cellular immune response. Comparing the mean gene expression profiles across the three time points, we see that both infection and selection with parasitoids alter gene expression in the larval fat body. We found that when flies adapted to high rates of parasitism, the uninfected larval fat body exhibits similar transcriptional changes as the fat body of infected flies that have not undergone parasitism every generation. To avoid ascertainment bias, we used data from an independent study of the larval fat body to define a set of 329 parasitism-responsive genes that were significantly differentially expressed with a log_2_ fold change (log_2_FC) of greater than one after parasitoid infection.

Among these genes, the constitutive changes in expression in the populations adapted to high rates of parasitism were correlated with the infection-induced expression in the populations that were not evolved with parasite pressure (Figure 2A). However, the transcriptional response seen after infection was greater than the change in gene expression seen in uninfected larvae that had evolved resistance (Figure 2A). This correlation in transcriptional profiles shows that the humoral immunity was partially activated in the evolved populations before the larvae were infected.

**Figure 2.**
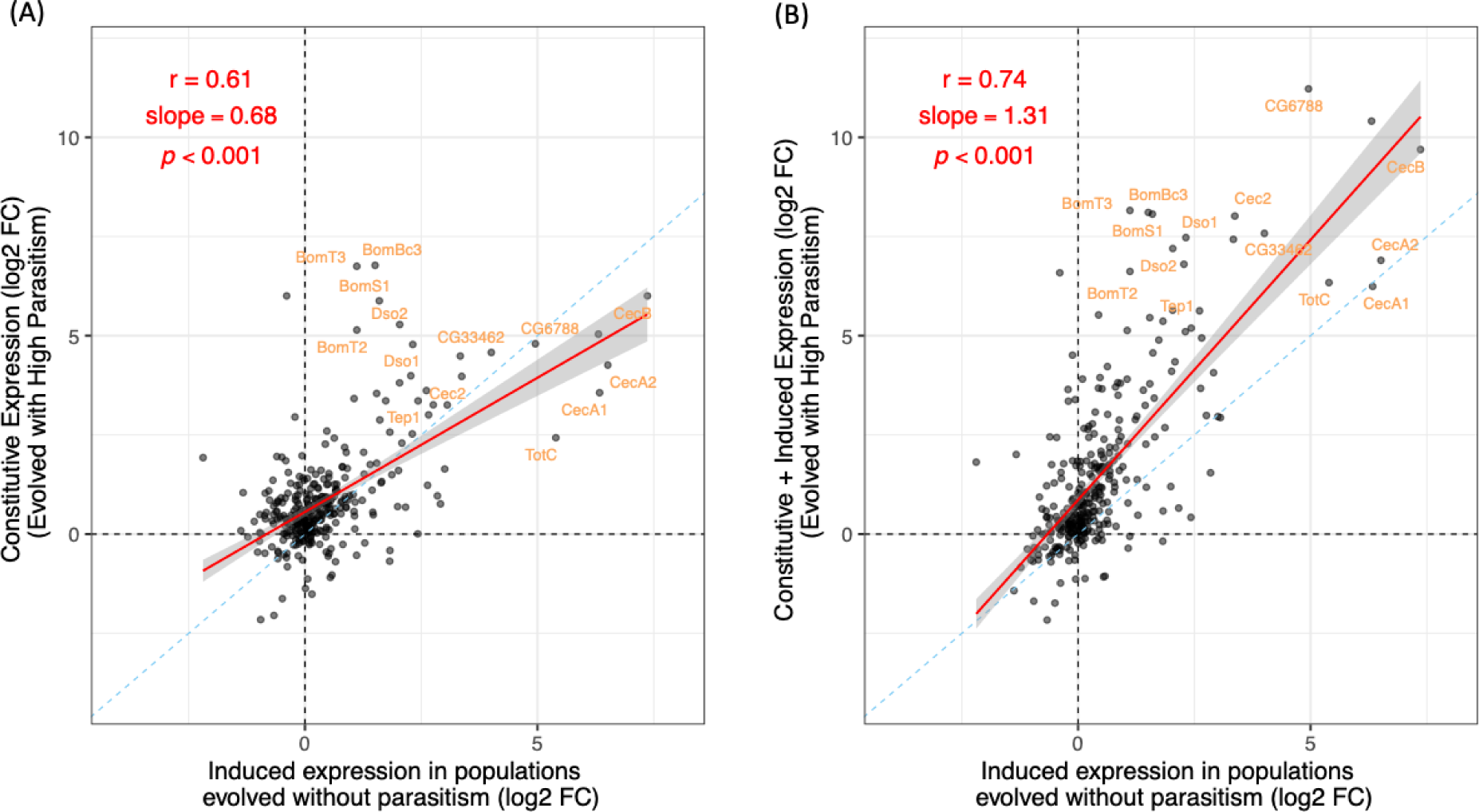
Changes in gene expression in the fat body following selection for resistance and parasitoid infection. In both panels the X axis represents the change in gene expression following infection in populations maintained without parasitoid infection. (A) Y axis represents the constitutive (uninfected) gene expression in populations selected for resistance to parasitoid *L. boulardi* infections relative to control populations. (B) Y axis represents the combined induced and constitutive changes in gene expression in populations selected for resistance to parasitoids, which is the expression profile of the selected populations under infected conditions in comparison to the control populations under uninfected conditions. The dotted blue diagonals indicate the 1:1 line. Red lines indicate fitted linear models to the data, with shaded area as 95% confidence intervals. *r* = Pearson correlation coefficient. Relative expression as log_2_FC.

The genes that show large transcriptional changes with constitutive induction or after infection tend to be upregulated as opposed to downregulated, which is consistent with the role of the fat body in secreting immune effectors (Figure 2). Among the 33 genes that showed log_2_FC of greater than 4 with constitutive induction and/or infection (Figure 2B), 73% are predicted to be secreted, compared to 9% in the full set of genes included in the RNAseq. This indicates that the genes showing large transcriptional inductions in the fat body are significantly enriched for secreted factors (Fisher’s Exact Test, *p*-value =1.99 ×10^−19^_)._

To further understand the effect of selection on the basal humoral immunity level of the selected populations, we compared the expression profiles between selected and control populations, under uninfected conditions across the three time points. After filtering out lowly expressed genes, we detected 9962 genes in the fat body. There is a total of 338 significantly differentially expressed genes with absolute log_2_FC of greater than 1. Taking the 30 most significant of these, the pattern of selection increasing expression more than decreasing expression is apparent (Figure 3A). These highly significant genes include short secreted peptides of the Bomanin family, which are regulated by the Toll pathway with antifungal and antibacterial activities [20], are more highly expressed in selected populations (Figure 3A). Upon a gene ontology (GO) enrichment analysis of the upregulated genes, we also observed an enrichment of serine-type peptidase activity (GO: Molecular Function) in the populations evolved with high rates of parasitism (Figure S1A). For instance, *CG30090*, *CG33462*, and *SPH93* are all serine proteases (Figure 3A) [21]. Proteolytic cascades of extracellular serine proteases play an important role in the regulation of immune responses, including the melanization reaction and activation of the Toll pathway [17,18,22–24].

**Figure 3.**
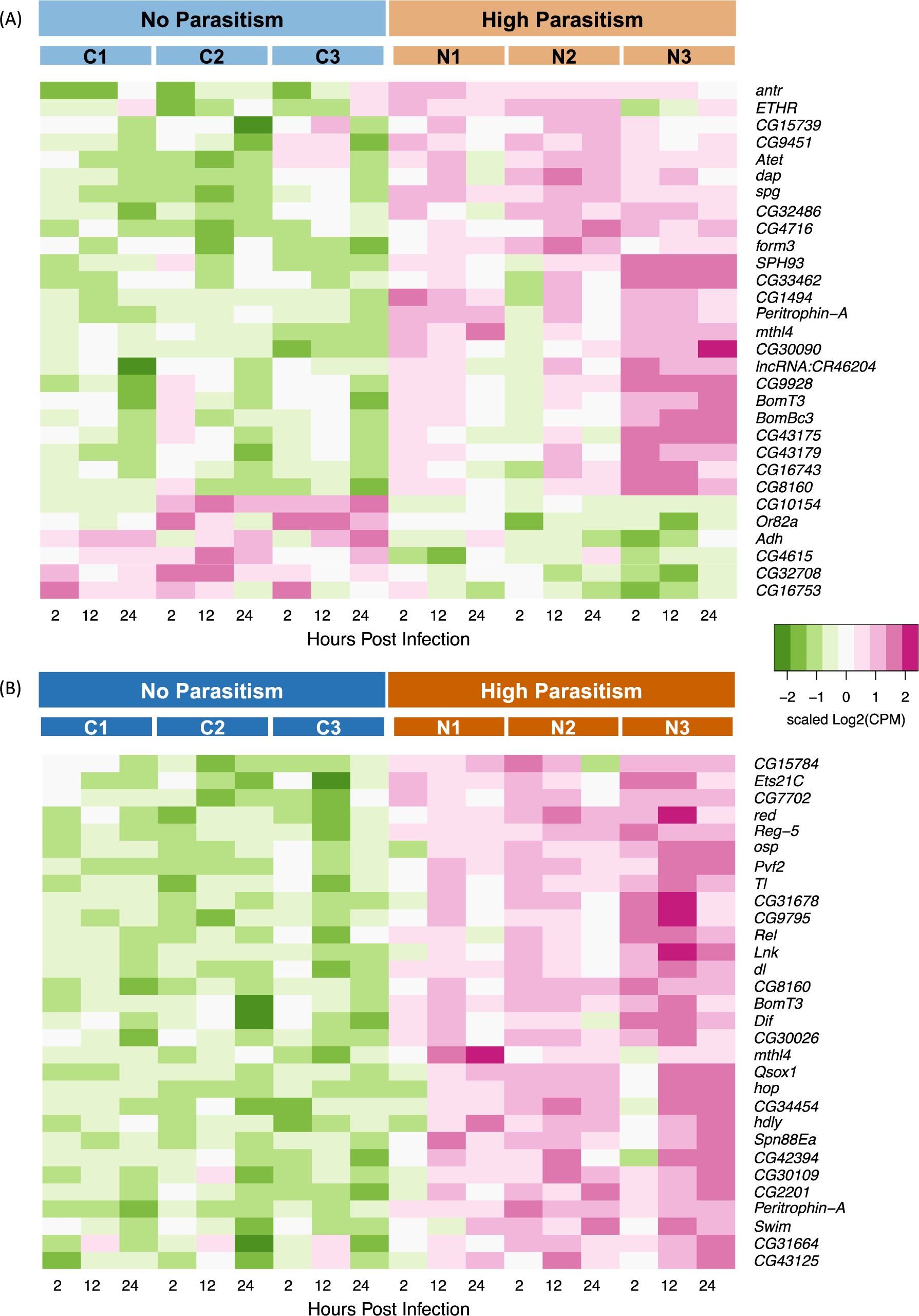
Differential expression of genes in the fat body between selection regimes. (A) The 30 most significantly differentially expressed genes under uninfected conditions over three time points (2, 12, 24 hpi). (B) The 30 most significantly differentially expressed genes after infection by the parasitoid wasp *L. boulardi* NSRef strain. Expression as scaled log_2_ counts per million (CPM).

### Populations evolved under the pressure of high parasitism rates can induce gene expression to a greater level

We see a higher level of humoral response in the evolved populations. Following infection, the parasitism-inducible genes showed an overall higher level of expression in the populations that had evolved under high rates of parasitism than those evolved without parasitism (Figure 2B). This was in contrast to what we previously found for cellular immunity, where the gene expression profiles become similar between the selection regimes after infection [8].

Under infected conditions, 398 significantly differentially expressed genes, with false discovery rate (FDR) of less than 0.05 and absolute log_2_FC of greater than 1, were found between the populations evolved with high rates of parasitism and those maintained without, indicating that there are still substantial differences in transcriptional expression even with parasitoid infection. The genes that are differentially expressed largely show higher expression in the populations evolved with high parasite pressure (Figure 3B). Strikingly, key components of three major immune signaling pathways (Toll, Imd, and JAK/STAT pathways) all show up within the top 30 significantly differentially expressed genes: *Tl*, *Dif*, *dl*, *Rel*, and *hop*. These genes are all significantly upregulated in the populations evolved with high parasitism, suggesting that these pathways are more active in the selected populations under the infected state. GO enrichment of significantly upregulated genes showed enrichment of terms related to “immune system process”, “defense response”, and “interspecies interaction” (Figure S1B). Therefore, the populations evolved with high parasite pressure displayed an overall more activated humoral immune response in the presence of parasitoid infection.

A number of parasitism-responsive genes showed similar levels of constitutive expression in the selected populations as the infection-induced expression in the control populations; these genes then became even more upregulated in the selected populations with infection (Figure 2). For example, *Tep1* is known to play a role in the melanization of parasitoid eggs [25], and it was expressed more highly in selected populations after infection. Similarly, the serine protease *CG33462* and the fibrinogen-like protein *CG6788*, in addition to a group of antibacterial cecropins, all showed further induction with infection in the populations evolved with high parasitism rates (Figure 2). It is also notable that virtually all the highly induced parasitism-responsive genes have greater levels of expression in the selected populations with infection than in the control populations (Figure 2B; log_2_FC>3 in control populations).

### The immune response is faster in populations evolved under strong parasite pressure

As the success of the anti-parasitoid immunity is dependent on how fast the fly can launch its defense [26], we hypothesized that the populations evolved under high rates of parasitism had an accelerated transcriptional response with infection. We analyzed the expression changes of 44 significantly differentially expressed genes with immune-related functions 2-, 12-, and 24-hours post-infection (hpi). There is generally higher upregulation of these immune-inducible genes at the 2 hpi time point in the populations maintained with parasitic pressure, relative to the baseline of uninfected populations maintained without parasitism (Figure 6). By 12 hpi, the level of induction of these genes became comparable between the populations. Interestingly, at 24 hpi, the populations evolved with parasitism again showed slightly higher level of transcriptional upregulation of these immune-inducible genes. This result suggests that the significantly differentially expressed immune-related genes are more rapidly and highly upregulated in the evolved populations at the early time point of 2 hpi.

### Only populations evolved under high parasitism rates activate the JAK/STAT pathway after infection

Parasitoid wasp infection can cause hemocytes to secrete the cytokines Upd2 and Upd3, leading to the activation of the JAK/STAT pathway in other tissues [19]. As we had observed upregulation of *hop* (which encodes the *Drosophila* JAK) in the populations adapted to high parasitism rates (Figure 3B), we investigated how JAK/STAT activity had evolved in these populations. We crossed males from the selected or control populations to females expressing a JAK/STAT pathway activity reporter [27]. The reporter expresses GFP under the control of ten Stat92E binding sites from the Stat92E-regulated gene *Socs36E*. At 24 hours post-infection (hpi), we see that JAK/STAT activity is strongly induced in the F_1_ progeny from the evolved populations, while there is little or no induction in those from the control populations (Figure 4; Welch’s *t*-test, d.f. = 2.95, *p*-value = 0.0238). Our results suggest that JAK/STAT is activated only after infection only in populations evolving under high parasitism pressure. The JAK/STAT pathway has been shown to be involved in the encapsulation response, where loss-of-function mutations in *hopscotch* results in reduced ability to generate lamellocytes and reduced encapsulation capacity [28]. Thus, the differential activation of JAK/STAT activity between the selected and control populations may play a part in the difference in resistance to parasitoid infection.

**Figure 4.**
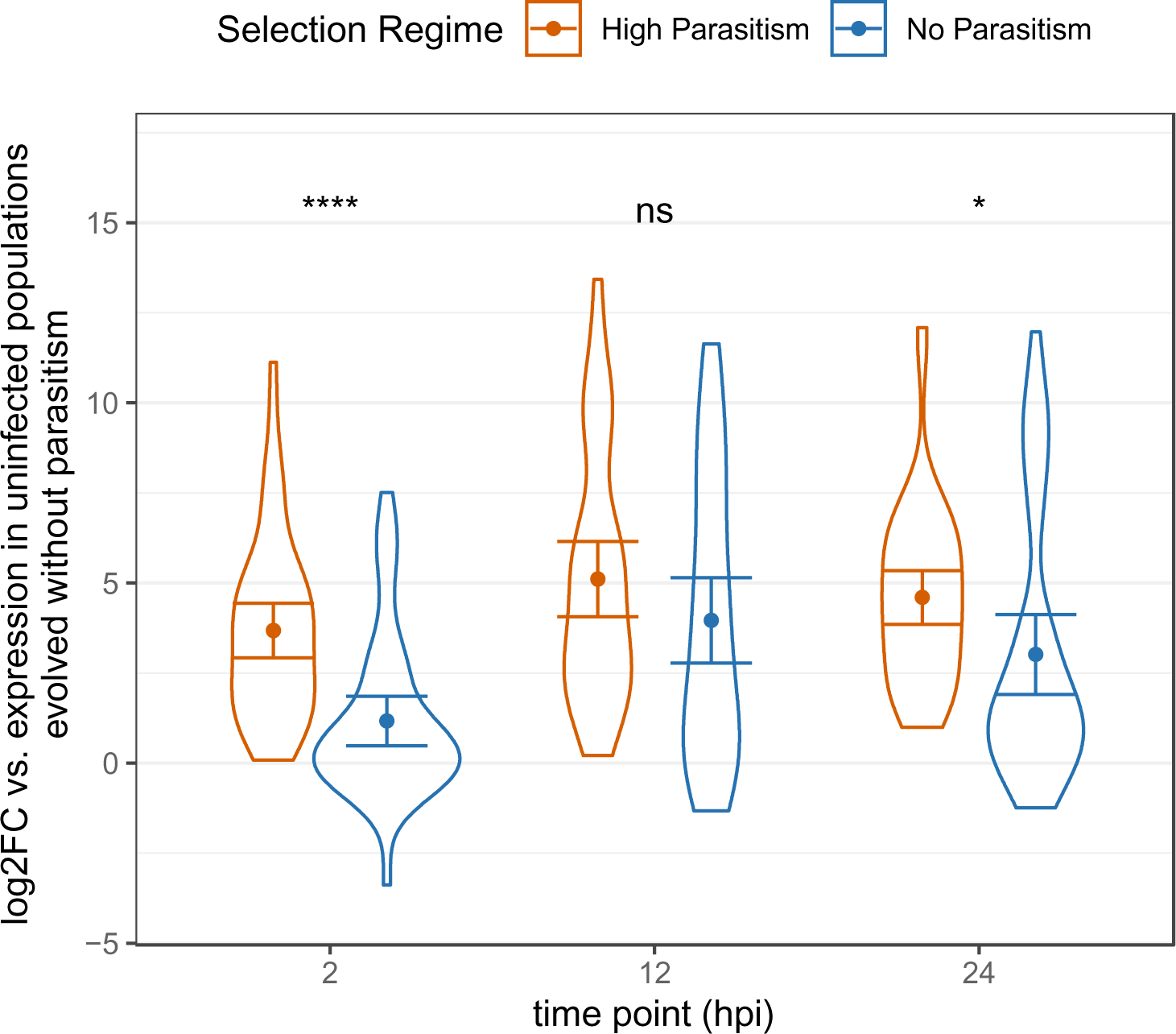
The speed of response following infection in the populations evolved with high parasitism and those maintained without parasitism. The induction of immune genes in the fat body is faster in populations evolved with high parasite pressure (at each time point, t-test, *df* = 86, **** *p* < 0.00005, * *p* < 0.05). Only genes with immune-related functions shown. Error bars representing 95% confidence intervals.

JAK/STAT activation in somatic muscles is the result of the cytokines Upd2 and Upd3 being secreted from hemocytes. In a previous study, we found that the immature lamellocyte cell states LAM1 and 2, which are typically only induced by parasitoid infection, become constitutively present in populations selected under high parasitism; the mature LAM3 lamellocyte cell state remains a largely inducible response [8]. To investigate whether this could underlie the difference in JAK/STAT activation, we examined the expression of Upd2 and Upd3 in the different hemocyte cell states present in a *D. melanogaster* larva [8,19]. Utilizing our previous single-cell RNAseq data [8], we found that LAM2 cells are the main *upd3-*expressing cell state, while plasmatocytes and crystal cells show almost no *upd3* expression (Figure 5A). The expression of *upd3* does not change markedly within a cell state upon infection. *upd2* is not included in this dataset as it did not pass the detection thresholds.

**Figure 5.**
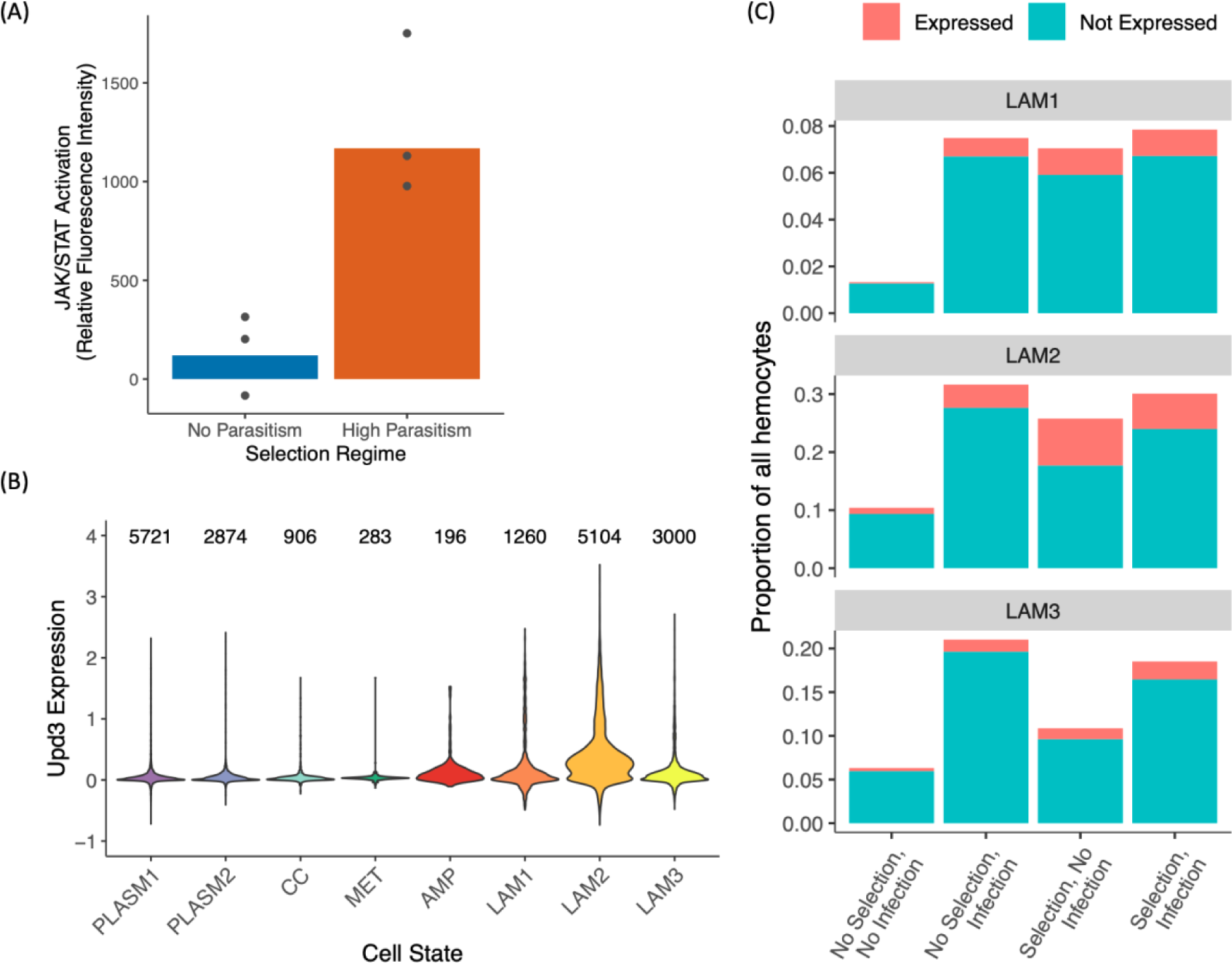
Differential JAK/STAT pathway activity between the selection regimes. (A) Females carrying a JAK/STAT pathway activity GFP reporter were crossed to males from the populations maintained under the different selection regimes. F_1_ progeny were assayed for GFP expression. Points represent replicate populations. At least 12 replicates for each condition were assayed for each population, where each replicate is a pool of 10 larvae. (B) Expression of *upd3* across the eight hemocyte cell states present in *D. melanogaster* larvae. Y axis represents normalized expression where feature count is divided by total count for each cell and scaled. (C) Proportion of different lamellocyte cell states after infection and selection, subdivided based on expression of *upd3*, where a cell is classified as *upd3*-expressing if the raw read count was above 0.

LAM2 cells are constitutively present in the populations evolved with high parasitism (Figure 5B), and their role as the cytokine-secreting hemocytes suggests that they might act as an early sensing system for the detection of wasp infection in the larvae. When we compare the expression of *upd3* in the three LAM cell states between the evolved and non-evolved populations, we see that in uninfected conditions, there is a higher proportion of *upd3-* expressing cells in the evolved populations compared to the controls across all three cell states (Figure 5B). Of the LAM2 cell state only, the evolved populations have 8-fold more cells expressing Upd3 than the control populations (Figure 5B). LAM2 is the final immature cell state before differentiation into the mature lamellocyte cell state LAM3 on the hemocyte lineage from plasmatocytes to lamellocytes [8]. Our results suggest that the immature LAM2 cells may play a role in the crosstalk between different parts of the immune system through cytokine secretion, by which they are likely responsible for the cell non-autonomous activation of the JAK/STAT pathway in other larval tissues including the somatic muscles [19].

## Discussion

We investigated how humoral immunity in *Drosophila melanogaster* populations evolved under high parasite pressure. This work complements our previous results on how the cellular immune response changed in response to high rates of parasitism. In contrast to hemocytes, where differences in gene expression largely reflect the differentiation of specialist cell types for the encapsulation response [8], RNA sequencing in the larval fat bodies is studying the activation of immune pathways and secretion of immune effectors. Our results showed that constitutive humoral immune mechanisms are favored when the probability of parasitization is high, consistent with theoretical predictions [2,29]. The selected *D. melanogaster* populations showed constitutive expression of immune-inducible genes in their larval fat bodies, indicating constitutive secretion of immune effectors for the anti-parasitoid response, even in the absence of infection.

As parasitoid virulence is high, theory predicts that the parasite will select for higher investment in both constitutive and induced defenses [2]. Our results are consistent with this hypothesis in that we see an increase in both forms of defenses in the populations evolved with high rates of parasitism. However, when we investigated the cellular immunity of these populations, we did not see the same greater overall response—the only differences were before infection, and after infection the populations of immune cells in the selected and control populations were similar [8]. While these findings may be affected by the timepoints we examined, it suggests that the two branches of the immune system may differ in their evolutionary dynamics.

Between the selection regimes, we observe a stark difference in the induced response when we investigated JAK/STAT pathway activity. With infection, we see strong activation of JAK/STAT activity in the selected populations whereas the controls showed no induction (Figure 4). JAK/STAT is an evolutionarily conserved signaling pathway with functions in immunity and defense. It may provide a link between the evolutionary changes we have observed in different tissues as JAK/STAT signaling involves cytokines secreted after infection, which then bind to receptors found on other cells, activating the pathway through an intracellular signaling cascade. The final effector of the pathway is the transcription factor STAT, which translocates when active to the nucleus and activates transcription of target genes. Wasp infection causes two cytokines to be secreted by hemocytes, Upd2 and Upd3, which then activate JAK/STAT activity in other tissues through ligand binding [19]. This non-cell autonomous activation of JAK/STAT signaling in the somatic muscles of *Drosophila* larvae is required for lamellocyte formation and encapsulation [19]. It may also influence transcriptional changes we observed in the fat body, as septic injury results in hemocyte-specific Upd3 cytokine secretion that activates JAK/STAT signaling in the fat body [30]. Similarly, the expression of the Tot family of stress response genes and the thioester-containing protein Tep1 are controlled by JAK/STAT signaling in the fat body [30–32]. We have previously found that these evolved populations showed constitutively active cellular immunity – particularly the constitutive presence of immature lamellocyte cell state LAM2 [8]. Reanalysing this data, we found that these cells are the main Upd3-expressing hemocyte subtype. It is therefore possible that it is these cells that are responsible for activating JAK/STAT in the evolved populations.

It is known that parasitoid wasps employ various strategies to block and evade host immunity [33–36]. One explanation of our results is therefore that the virulent NSRef strain of *L. boulardi* wasps may be suppressing JAK/STAT pathway activity in susceptible flies while the populations maintained under high parasite pressure have evolved to counter this suppression. If this is the case, the constitutive expression of Upd3 cytokines by hemocytes in these evolved populations in the absence of wasp infection might allow these flies to activate JAK/STAT signaling before the parasitoid venoms can sabotage this defense. Altogether, our results suggest a hypothesis by which constitutively producing previously inducible precursor cell states, the evolved populations can more effectively respond to parasitoid infections through a two-fold mechanism: 1) the constitutive presence of immature lamellocytes allows for the rapid production of mature lamellocytes upon infection, and 2) the cellular immune system is also “primed” to activate other parts of the immune response through the secretion of cytokines by LAM2 before they differentiate into mature lamellocytes. Interestingly, the transcription of Upd3 does not increase in LAM2 cells after infection, so the activation of JAK/STAT may rely on release of cytokines being controlled at the level of translation or secretion, as is common in vertebrates [37,38].

Autoimmune damage may be a price hosts have to pay for resistance to infection. This cost may be present both in the absence of infection, as with many autoimmune disorders, or during infection, in what is known as hypersensitivity reactions often linked to inflammation and complement activation caused by an immune response [39]. While we did not investigate self-damage, our results are consistent with both effects: we see both constitutive activation of immunity in the absence of an infection and overall greater immune activation after infection. In particular, aberrant cytokine signaling through the JAK/STAT pathway underlies many autoimmune disorders found in humans [40]. Our results show both greater Upd3 expression by hemocytes in the absence of infection and higher JAK/STAT activity after infection, indicating potentially high immunopathological costs to the hosts in the evolved populations.

To our knowledge, this is the first model in which the evolution of humoral immunity under high parasite pressure has been studied. Along with our previous findings on the cellular immunity of these selected populations [8], we show that constitutive immunity is favored in two different immune tissues in the *D. melanogaster* larvae when selected for resistance with high rates of parasitism. Future studies on different species and models will provide more insights to how general is this observation of natural selection driving evolution of constitutive defenses when infection is common. Our studies on evolution over short time scales may provide an explanation more broadly of the evolutionary logic as to why some aspects of host immunity are inducible and others are not, and the reasons why immunopathological disease is so widespread.

## Materials and Methods

### Artificial selection of *Drosophila melanogaster*

The *D. melanogaster* lines used were continuations of selections from the populations established by Leitão et al. [8], where the initial source populations were established using isofemale lines founded from 377 females collected in Cambridge, UK in July 2018 using banana and yeast traps set up in an allotment plot (52^°^12’12.5”N 0^°^09’00.6”E). Isofemale lines were established by placing single females in vials with cornmeal food (per 1200 ml water: 13g agar, 105g dextrose, 105g maize, 23g yeast, 35ml Nipagin 10% w/v). Five females and five males were collected from the progeny of each isofemale line to create a source population of 3770 flies. The source population was collected into cages and fitted with 90mm apple agar plates (per 1500 ml water: 45g agar, 50g dextrose, 500ml apple juice, 30ml Nipagin 10% w/v) covered with yeast paste (*Saccharomyces cerevisiae* – Sigma-Aldrich #YSC2). The flies carried out overnight egg lays and eggs were collected from the agar plate with phosphate-buffered saline (PBS) using a paintbrush. The eggs were collected in 15 ml centrifuge tubes and allowed to settle to the bottom. Subsequently, 500 µl of the egg solution was transferred into a 1.5ml microcentrifuge tube, from which 6 µl of egg solution was added to plastic vials containing cornmeal food. The vials were kept at 25°C, in a 14-hr light/10-hr dark cycle and 70% humidity. 48 hours after egg transfer, a single female wasp was added into each vial for infection of 24 hours, and some vials were not infected. Vials were then incubated at 25°C for 12 days in total, then flies from infected treatments were collected and randomly sorted into triplicate selection lines (N1-3). Flies that were not infected were sorted into triplicated control populations (C1-3). Each subsequent generation of selected populations were maintained in the same way, while the control populations were maintained with the same protocol without infections. The population sizes were maintained at around 200 adult flies for each.

### Encapsulation assays

Female wasps lay eggs inside the *Drosophila* larval hemocoel through the ovipositor. If successful, the wasp larvae would feed on *Drosophila* larval tissue and emerge from the pupae of the fly as the adult wasp. If the *Drosophila* is successful in its immune defense, the flies encapsulate the wasp eggs, resulting in black melanized capsules which can be visualized under a microscope.

Larval density was controlled for the encapsulation assays in a similar way as the artificial selections (see above section). Two days after egg transfer into cornmeal vials, three female wasps of the *Leptopilina boulardi* strain NSRef were added to each vial for 3 hours of infection at 25°C. For each fly line, we prepared 10 vials with no infection and 20 vials with infection. We then counted the flies that emerged from the vials without infection. Flies that emerged from the infection vials were checked for the presence/absence of encapsulation capsules by squishing anaesthetized flies between two glass microscope slides. These flies were then observed under a dissecting microscope and counted. The formula for calculating encapsulation ratio is as follows:

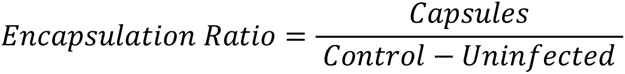

Capsules is the mean number of flies from infection vials that have visible capsules and Uninfected is the mean number of flies from infection vials with no discernable capsule, and Control is the mean number of flies emerging from vials with no infection. We can then estimate the proportion of infected flies with successful encapsulation per line.

### Wasp maintenance

*Leptopilina boulardi* strain NSRef [41] was maintained using a susceptible *D. melanogaster* outbred population. Eggs were added to vials of cornmeal fly food as described above. Two female wasps and one male wasp were added to each vial, where the vials were then incubated at 25°C for 24 days, in a 14-hr light/10-hr dark cycle and 70% humidity. Adult wasps were collected and maintained in cornmeal vials with a drop of honey added to the cotton plug.

### Larval fat body preparation

Larval fat bodies were dissected from late 2^nd^ instar to early 3^rd^ instar *D. melanogaster* larvae. The whole fat body was obtained by first removing the head of the larva then pulling back the cuticle to expose the fat body and other inner organs. The gut, salivary gland, and other organs were then removed and discarded. Each fat body is dipped in a clean drop of PBS immediately following dissection to wash off any hemocytes that might be attached to the surface, and then transferred into a screw cap 0.5 ml tube containing 50 µl of ice-cold PBS. After 10 fat bodies had been transferred and pooled in a tube on ice, the fat bodies were then spun down by pulse centrifugation. The PBS supernatant was then removed by pipetting carefully off the top without disturbing the fat bodies which had collected at the bottom of the tube.

### Library preparation for RNA sequencing

At 2 hpi, 12 hpi, and 24 hpi, groups of 10 male larvae were dissected for fat bodies from each of the control and selected populations. Fat bodies from uninfected larvae were also dissected from groups of 10 male larvae from each population at the same time points. RNA was isolated from the dissected fat body tissues. For each sample, as soon as 10 fat bodies were pooled, 350 µl of Tri-reagent (Ambion 10296010) was added immediately, and the fat bodies were homogenized by vortexing for 5-10 seconds. 70 µl of chloroform was added and the tubes were shaken for 15 seconds and then incubated at room temperature for 3 minutes. Samples were then centrifuged for 10 minutes at 12,000g at 4°C. 100ul of the upper aqueous phase was transferred to a fresh Eppendorf tube and 175 µl of propan-2-ol was added and mixed by inverting the tubes several times. After incubating 10 minutes at room temperature, the tubes were centrifuged at 12,000g at 4°C for 10 minutes. The supernatant was removed and 350 µl ice-cold 70% ethanol was added. Centrifuge at 4°C for 2 minutes at 12,000g and the ethanol was removed and the RNA pellets were air dried briefly and 20 µl of nuclease-free water was added. The RNA pellets were fully dissolved by incubating tubes at 45°C for 5 minutes on a heat block. RNA was quantified using a Qubit RNA HS assay kit (Thermofisher Q32852). RNAseq libraries were prepared using a NEBNext Ultra II Directional RNA Library Prep Kit for Illumina (New England Biolabs E7760S) with NEBNext Multiplex Oligos for Illumina (96 Unique Dual Index Primer Pairs) (New England Biolabs E6440S) and polyA enrichment module (NEB E7490S). Up to 1µg of total RNA was used to make each library. mRNA was fragmented at 94°C for 15 minutes after polyA enrichment, according to the manufacturer’s recommendations. Adaptor concentrations and number of amplification cycles were adjusted according to the amount of starting material. The quantity of each prepared library was then measured using Qubit DNA HS assay kit (Thermofisher Q32851). The quality of each library was assessed using a Bioanalyzer high sensitivity kit on an Agilent 2100 Bioanalyzer (Agilent 5067-4626). The libraries were submitted for sequencing at Cancer Research UK Cambridge Institute Genomic Core Facility using Illumina NovaSeq with 100bp single-end reads.

### Statistical analyses of RNAseq data

Trim Galore (https://www.bioinformatics.babraham.ac.uk/projects/trim_galore/) was used to trim raw RNAseq reads, with a Phred score of 20 for quality control. Reads with fewer than 50 bases after trimming were removed. The reads were then mapped and counted using STAR v2.6.0 [42] to *D. melanogaster* reference genome Dmel-r6.43.

The R package edgeR (RRID:SCR_012802) was used for differential expression analyses. Genes that had a count per million (CPM) above ten in at least three libraries were kept. This served as a threshold for identifying genes with detectable expression in our dataset. To increase our statistical power, we combined the data across the three time points in our analysis and compared the mean gene expression profiles. Dispersions were estimated using the Cox-Reid profile-adjusted likelihood (CR) method in edgeR, and the expression data were fitted with a negative binomial generalized linear model (GLM). Differential expression of genes was then determined with a quasi-likelihood (QL) *F*-test, where a false discovery rate (FDR) of 0.05 was set as the significance threshold.

When comparing expression levels between selected and control populations, we combined the data across the three time points at which the samples were collected and only contrasted between selection and immune challenge.

### Changes in gene expression following selection/infection analysis

Data from an independent study of the *D. melanogaster* larval fat body were used to define a set of 329 parasitism-responsive genes that were significantly differentially expressed with a logFC of greater than 1 following wasp parasitization. The raw RNAseq reads from this independent study were deposited in the NCBI Sequence Read Archive under the BioProject number PRJNA1021619.

### Enrichment for secreted proteins analysis

A list of all secreted proteins in *D. melanogaster* was extracted from *Drosophila melanogaster* extracellular domain database (FlyXCDB) [43].

### Counts-per-million (CPM) expression heatmap analysis

Expression data were produced using the R package edgeR, where moderated log_2_ counts-per-million (logCPM) data were computed. The logCPM values were then scaled for each gene in the heatmaps, by centering on the mean value and dividing by standard deviation. The gplots and RColorBrewer (RRID:SCR_016697) packages was used for visualization of the data.

### Speed of response analysis

To analyze the differential speed of anti-parasitoid response between the populations selected under high rates of parasitism and the control populations, we first extracted a list of genes that are significantly upregulated with logFC over 1.5 for either the populations adapted to high rates of parasitism or the control populations, averaging the expression profiles over the three time points included in the RNAseq. Within this list of significantly upregulated genes, we then extracted immune-related genes corresponding to the FlyBase GO annotation “Immune System Process” (GO:0002376) and all its daughter terms.

The logFC of each of these genes was then computed against a baseline expression level of the control populations under uninfected condition at each time point. A two-sample t-test was used to compare the mean logFC at each time between selected and control populations.

### Gene Ontology enrichment analyses

A Gene Ontology (GO) enrichment analysis was conducted using the R package goseq [44]. The list of genes detected in RNAseq analysis was used as a background set. Sets of significantly upregulated and significantly downregulated genes with an FDR < 0.05 and up or down-regulation of at least one logFC were then analyzed separately to determine GO enrichment. The length bias inherent to RNAseq data were accounted for by calculating a Probability Weighting Function (PWF) using goseq, that gives a probability that a gene will be differentially expressed based on its length alone. A null distribution for GO category membership was approximated with Wallenius distribution, and each GO category is then tested for over and under representation amongst the set of DEGs.

Where there are more than 10 GO terms in any of the three major Gene Ontology branches (Biological Processes, Cellular Components, and Molecular Functions), the list of GO terms was summarized using REVIGO [45] (RRID:SCR_005825).

### JAK/STAT pathway activity reporter assay

JAK/STAT pathway activity was evaluated by preparing samples of larval tissue lysate. At least eight independent samples were made from each cross (between females carrying reporter construct and males from each selection regime). Eight to ten third instar larvae are collected for each sample, and tissuelysed in 100ul of PBS with about ten 1.0mm diameter zirconia/silica beads (Thistle Scientific # 11079110z) using Qiagen TissueLyser II (Qiagen # 85300) for 2 minutes at 30Hz. The samples were immediately spun down at 4000rpm for 10 minutes at 4°C, and 50µl of the supernatant for each sample were transferred into a well in a flat clear-bottom black polystyrene 96-well plate (Corning # 3603) for fluorescence reading with a SpectraMax iD3 Plate Reader (Molecular Devices) using the SoftMax Pro 7 software. GFP fluorescence intensity is measured with excitation at 485nm and emission at 535nm. The rest of the supernatant for each sample is used to measure the total protein level of the sample with a Bradford assay (Merck # B6916), following reagent protocol. The total protein level was used as a normalization of GFP relative intensity.

### JAK/STAT activity analysis

We tested for the difference in the induction of JAK/STAT activity between the selection regimes using the method described above and analyzed the data with a linear mixed-effects model using the function below using the R package nlme [46]:

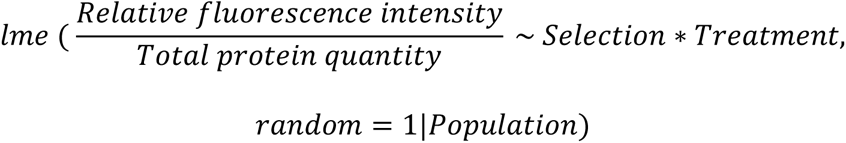

### Single-cell RNA sequencing analysis

The single-cell RNAseq analysis was done using our previous dataset from Leitao et al. [8]. Analysis was carried out using the R packages Seurat [47] (RRID: SCR_007322) and reshape2 [48].

## Data availability

The RNAseq data has been submitted to the NCBI Sequence Read Archive under the BioProject number PRJNA102111, with BioSample accessions from SAMN37513352 to SAMN37513387. Processed data files and scripts to analyze data are available on Apollo, the University of Cambridge repository (https://doi.org/10.17863/CAM.101651).

